# A developmentally-inspired hypoxia condition promotes kidney organoid differentiation from human pluripotent stem cells

**DOI:** 10.1101/2023.07.29.551084

**Authors:** Hyeonji Lim, Dohui Kim, Haejin Yoon, Joo H. Kang, Dong Sung Kim, Tae-Eun Park

## Abstract

Kidney organoids derived from human pluripotent stem cells (hPSCs) lack a contiguous network of collecting ducts, which limits their utility in modeling kidney development and disease. Here, we report the generation of kidney organoids containing ureteric bud (UB)-derived collecting ducts connected to metanephric mesenchyme (MM)-derived nephrons using developmentally-inspired hypoxic differentiation conditions. Hypoxia promotes a reiterative process of branching morphogenesis and nephron induction through reciprocal interactions between co-induced MM and UB, which lead to a higher-order kidney organogenesis *in vitro*. The resulting kidney organoids demonstrate greater maturity, as indicated by higher levels of functional markers and more realistic micro-anatomy of the tubules and collecting ducts. Additionally, these hypoxic-enhanced kidney organoids show a great potential as *in vitro* models for renal cystic diseases, as they efficiently generate cystic formations and display high sensitivity to drugs. This hypoxia approach may open new avenues for an enhanced understanding of kidney development and diseases.

## INTRODUCTION

During kidney development, intricate and elaborate structures are formed by reciprocal interactions between the ureteric bud (UB) derived from the anterior intermediate mesoderm (aIM) and the surrounding metanephric mesenchyme (MM) derived from the posterior IM (pIM).[1] Signals from the MM induce the UB to branch repeatedly, giving rise to the entire network of collecting duct, and simultaneously, signals from the UB induce the MM to form nephrons.[2, 3] Therefore, the generation of UB and MM is crucial for reconstituting a complete kidney structure *in vitro*, which has nephrons connected by collecting ducts that finally converge into the ureter.

Unfortunately, most protocols related to kidney organoids have focused on generating MM-derived nephron progenitor cell (NPC) lineages, such as podocytes, Bowman’s capsules, and renal tubules, thereby limiting the induction of UB-derived ureteric bud progenitor cells (UPCs).[4–8] The method is based on the knowledge that pIM migrates out of the primitive streak (PS) later than does aIM,[9] thereby being exposed to long WNT signaling in the primitive streak.[10] For example, the Morizane protocol generate late PS from human pluripotent stem cells (hPSCs) through long-term WNT activation with CHIR99021 (CHIR) and differentiation to pIM with Activin A followed by the maturation of MM with fibroblast growth factor 9 (FGF9) to induce the formation of NPCs.[6] Although this selective induction of NPCs enables efficient generation of the complex structure of human nephrons *in vitro*, the formed nephrons are separately positioned within the organoids, and their organization and function do not mimic those of a mammalian kidney because of the lack of UB-derived cells and signals. The Takasato protocol generates both MM- and UB-derived cells by controlling the exposure time to Wnt signaling, FGF9, and retinoic acid; however, UB-like cells resemble NPC-derived distal renal tubules rather than the collecting duct system.[4] Although these nephron organoids have been used to model kidney diseases, such as polycystic kidney diseases (PKDs),[8, 11–16] their disease models do not consider the pathogenesis of collecting duct origins, which limits the potential for discovering broad-spectrum drugs.

Given the role of reciprocal interactions between UB and MM during kidney organogenesis, a promising strategy for recreating a high-order kidney structures *in vitro* is to generate UB from primary mouse/rat tissue or hPSCs and assemble them with NPCs. They have showed robust UB branching along with differentiated nephrons located at the UB tips, similar to the kidneys *in vivo*. However, this strategy requires animal-derived stromal cells for the maturation of organoids,[17] the resulting assembloids lack vascular structures,[17–19] and more laborious.

In this study, we aimed to differentiate the human induced pluripotent stem cells (hiPSCs) into UB- and MM-lineage cells in the same culture dish to enhance the structural and functional maturation of kidney organoids. Our approach simulates oxygen tension during kidney development, which results in generation of kidney organoids containing a UB-derived collecting duct tree connected to NPC-derived proximal, intermediate, and distal tubules, similar to the *in vivo* kidney structure. We demonstrated that the regulated reciprocal signals between UB and MM in early stage of differentiation were responsible for the development of entire renal branching patterns, tubular elongation and improved maturity. Furthermore, we successfully modelled PKDs in hypoxia-enhanced kidney organoids, which exhibited cyst formation in the entire tubular structure with excellent sensitivity to drugs against PKDs.

## RESULTS

### Hypoxic stimulation promotes commitment to UB and MM lineages

Hypoxic stimulation can promote the commitment of progenitors to various cell types, and microenvironmental oxygen concentration is an important signal for regulating embryo development.[20–22] To explore the possibility that oxygen regulation could improve the formation of organized kidney structures, we modified the Morizane protocol[6] by shifting the culture to hypoxic conditions (5% O_2_) for 9, 14, and 21 days, corresponding to NPC, renal vesicle, and nephron stages, respectively (Figure S1). Kidney organoid differentiation under 5% O_2_ for 9 days and 21% O_2_ for the next 12 days resulted in the highest induction of tubular structures, which showed distinct morphological differences from organoids cultured according to the original Morizane protocol (‘Normoxia’) (Figure 1A and Figure S1). Cells exposed to 1% instead of 5% O_2_ for 9 days did not produce self-organized renal structures (Fig. S1), suggesting that oxygen can stimulate kidney differentiation under precise levels and temporal dynamics of oxygen tension.[23] We further investigated whether the optimized hypoxic condition (‘Hypoxia’; 5% O_2_ for 9 days) stimulated the differentiation of hiPSCs into UB and MM (Figure 1A).

**Figure 1.**
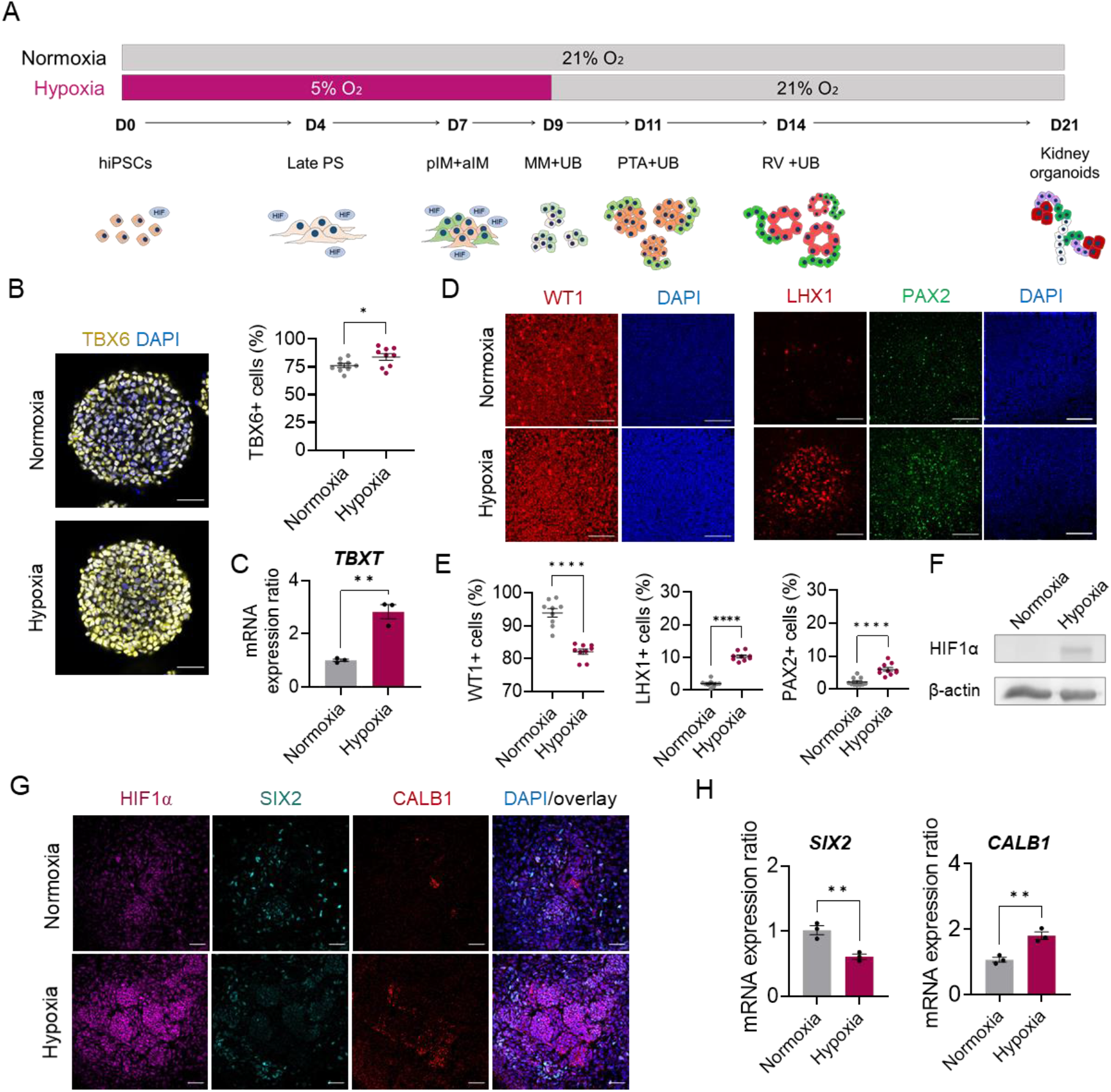
Dual induction of MM and UB from hiPSCs under hypoxic stimulation. (A) Timeline for *in vitro* differentiation of the kidney organoids from hiPSCs under normoxic and hypoxic conditions. hiPSCs, human induced pluripotent stem cells; PS, primitive streak; pIM, posterior intermediate mesoderm; aIM, anterior intermediate mesoderm; MM, metanephric mesenchyme; UB, ureteric bud; PTA, pre-tubular aggregate; RV, renal vesicle. (B) Left: Immunofluorescent micrographs at day 4 labelled with TBX6 (yellow). Scale bars, 50 µm. Right: quantification of the percentage of TBX6+ cells on day 4 under normoxic and hypoxic conditions analyzed by ImageJ software. (C) qRT-PCR on day 4 showing the expression of marker of PS (*TBXT*) in cells cultured under hypoxic versus normoxic conditions. (D) Immunocytochemistry for pIM marker, WT1, and aIM markers, PAX2 and LHX1, on day 7 of differentiation. Scale bars, 100 µm. (E) Percentage of cells positive for WT1, PAX2, or LHX1 analyzed by Image J. (F) Western blot showing HIF1α expression on day 1 under normoxic and hypoxic conditions. (G) Immunocytochemistry for HIF1α, SIX2 (MM marker), and CALB1 (UB marker) on day 9 of differentiation under normoxic and hypoxic conditions. Scale bars, 50 µm. (H) qRT-PCR on day 9 showing relative expression of *SIX2* and *CALB1* under hypoxic and normoxic conditions. All data are plotted as mean ± S.E. and n = 3 for the independent experiments. *P* values were determined by two-tailed unpaired t-test (\**P* < 0.05; \*\**P* < 0.001; \*\*\**P* < 0.0001; \*\*\*\**P* < 0.00001).

Following the Morizane protocol, we treated cells with CHIR and Noggin as a WNT activator and a BMP inhibitor for four days to induce TBX6+/TBXT+ PS formation under normoxic and hypoxic conditions.[6] Hypoxic conditions robustly stimulated PS induction, exhibiting 84% efficiency, whereas cells differentiated under normal conditions were 76% TBX6-positive (Figure 1B). A statistically significant increase in mRNA expression of *TBXT* was observed in cells on day 4 under hypoxic versus normoxic conditions (Figure 1C). This is consistent with recent findings that early mesoderm and endoderm-instructive genes, including TBXT, are selectively upregulated in PSCs under hypoxia, and that this regulation is mediated by hypoxia-inducible transcription factor 1α (HIF1α).[24, 25]

Subsequent treatment with Activin A between days 4 and 7 induced pIM cells (WT1+) as demonstrated in a previous study.[6] Under the hypoxic condition, however, proportion of pIM cells (WT1+) was lower than that under the normoxia condition (82.29% versus 93.65%). On the other hand, the percentage the aIM cells (PAX2+ or LHX1+) was significantly increased (Figure 1D, E), indicating that hypoxia facilitated the differentiation of hiPSCs into aIM cells, which is the origin of UB. The past *in vivo* and *in vitro* studies confirmed that HIF1α is activated in developing kidney, playing regulatory roles in nephrogenesis.[23, 26, 27] Indeed, western blot analysis revealed higher level of HIF1α when cells were exposed to the hypoxic condition during differentiation (Figure 1F). Confocal microscopic analysis further confirmed higher accumulation of HIF1α in hypoxic stimulated cells until day 9 (Figure 1G). Treatment with FGF9 between days 7 and 9 induces the differentiation of pIM into MM in the Morizane protocol.[6] Although the expression of SIX2, a MM marker was revealed on day 9 in both normoxic and hypoxic groups, the percentage of cells expressing SIX2 appeared lower when cultured in hypoxia (Fig. 1G), due to the decreased portion of pIM cells. Consistently, quantitative reverse transcriptase PCR (qRT-PCR) confirmed decreased mRNA expression level of *SIX2* in hypoxic condition (Figure 1H). Remarkably, we observed a significantly larger area of the UB cells by staining with CALB1 under the hypoxic condition than that under the normoxic condition, albeit without the formation of a CALB1+ duct-like structure. In line with the immunostaining result, higher mRNA level of *CALB1* expression was observed. Because MM and UB are derived from pIM and aIM, respectively, no overlap between SIX2+ and CALB1+ cells was noticed (Figure 1G, H).

### Developmentally-inspired kidney organoids exhibit elongated tubular structures including collecting ducts

We investigated the effects of dual induction of MM and UB under the hypoxic condition on the formation of high-order kidney organoid structures. On day 21, the resulting kidney organoids exhibited uniquely elongated tubular structures when exposed to 5% O_2_ for the first 9 days, whereas those generated without oxygen regulation had a typical nephron structure shown in the past study[6] (Figure 2A,B). When counting the number of kidney organoids larger than 200 µm, we found that the moderate hypoxic condition (5% O_2_ for 9 days) led to a higher efficiency of organoid formation than did the normal oxygen conditions (Figure 2C). Confocal microscopic analysis revealed the formation of glomerular podocytes (PODXL+), proximal tubules (LTL+), and distal tubules/collecting ducts (CDH1+) under normoxic and hypoxic conditions (Figure 2D). The area of glomerular region decreased by more than half (9.98% vs. 24.84%) under the hypoxic condition, whereas the area of the tubules/collecting ducts significantly increased (90.02% vs. 75.15%) (Figure 2E). The proximal and distal segment accounted 38.66% and 51.36% in hypoxia, and each was 4.54% and 10.33% of increase compared to the normoxic condition (proximal; 34.12%, distal; 41.03%) (Figure 2E). Striking increase of CDH1+ regions indicate that kidney organoids generated under regulated oxygen tension had developed tubular structures. Despite the decreased area of podocytes under hypoxic conditions, the podocytes had a more mature feature (figure S2A), as evidenced by the basally localized expression of ZO-1 (tight junction) observed in slit diaphragms in kidney tissue.[28]

**Figure 2.**
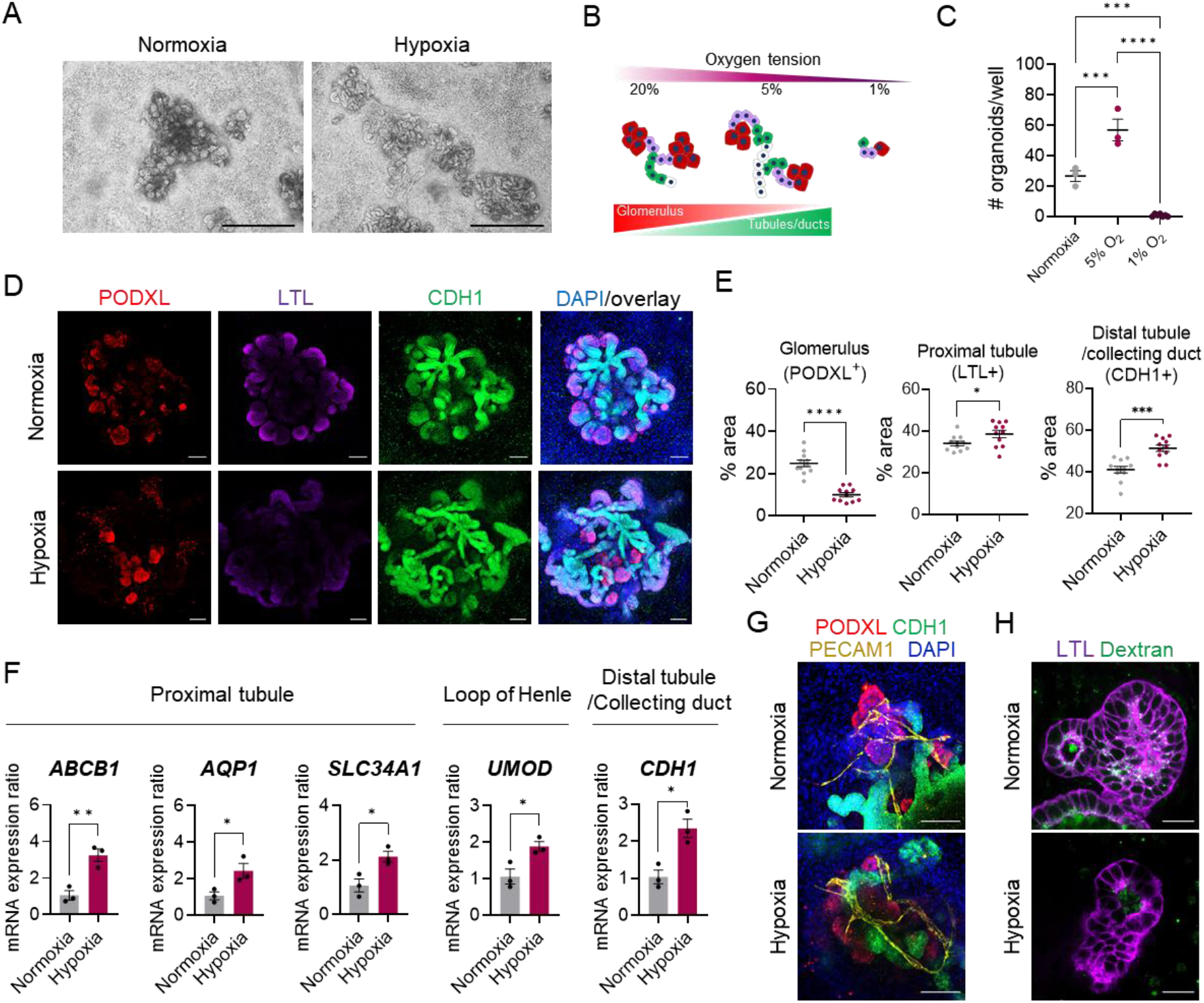
Maturation of kidney organoids with elongated and functional tubules under optimal hypoxia. (A) Bright-field images of kidney organoids on day 21 of differentiation under normoxic and hypoxic conditions. Scale bars, 500 µm. (B) Diagrams of different composition of nephron segments in the resulting kidney organoids depending on oxygen tension during initial nine days. (C) The number of kidney organoids > 200 µm per well under normoxia, 5% O_2_, and 1% O_2_ conditions on day 21. *P* values were determined by one-way ANOVA followed by Tukey’s multiple comparison test (\*\*\**P* < 0.0001; \*\*\*\**P* < 0.00001). (D) Immunocytochemistry for a marker of podocytes (PODXL), proximal tubules (LTL), and distal tubules/collecting ducts (CDH1) on day 21 of differentiation under normoxic and hypoxic conditions. Scale bars, 100 µm. (E) Percentage of cells positive for PODXL, LTL, or CDH1 on day 21 differentiated under normoxic and hypoxic conditions. (F) Graph showing the ratios of mRNA expression of genes encoding markers for proximal tubules (*ABCB1*, *AQP1*, and *SLC34A1*), loops of Henle (*UMOD*), and distal tubules/collecting ducts (*CDH1*) in the kidney organoids on day 21 differentiated under hypoxic relative to normoxic conditions. (E and F) *P* values were determined by two-tailed unpaired t-test (\**P* < 0.05; \*\**P* < 0.001; \*\*\**P* < 0.0001; \*\*\*\**P* < 0.00001). (G) Immunocytochemistry for PODXL, PECAM1, and CDH1 in kidney organoids on day 21 differentiated under normoxic and hypoxic conditions. Scale bars, 100 µm. (H) Dextran uptake assay showing endocytic uptake of LTL+ proximal tubules. Scale bars, 50 µm.

qRT-PCR analysis also revealed upregulated expression of NPHS1 (figure S2B), which functions in the glomerular filtration barrier.[29, 30] These findings suggest that the hypoxic condition do not negatively affect glomerular formation but rather stimulate glomerular cell maturation.[31] In line with the tubular elongation shown in the fluorescent images, the mRNA expression of *ABCB1*, *AQP1*, *SLC34A1* (proximal tubule), *UMOD* (loop of Henle), and *CDH1* (distal tubule/collecting duct) significantly increased under hypoxic conditions (Figure 2F).

Our study sought to determine whether the optimized hypoxic condition (5% O_2_ for 9 days) could enhance vascularization in kidney organoids, based on earlier findings that different hypoxic condition (7% O_2_) at later stages of kidney organoid differentiation increase the endothelial population.[32] We observed significantly higher mRNA expression of *PECAM1* in hypoxia-enhanced kidney organoids than in control organoids (Figure S2C); however, no significant change was noticed in the level of renal vascularization (PECAM1+) (Figure 2G). This finding suggests that the hypoxic condition is not optimal for enhancing kidney organoid vascularization. To examine whether hypoxia-enhanced kidney organoids exhibit physiologically relevant functions, we conducted an *in vitro* dextran uptake assay. After 24 h of treatment with 10 kDa dextran, kidney organoids could specifically take up and retain dextran within LTL+ proximal tubular epithelial cells (Figure 2H), indicating that kidney organoids were capable of reabsorption.

After examining whether UB was successfully differentiated into collecting ducts in hypoxia-enhanced kidney organoids, confocal microscopic analysis revealed that several nephrons stained with anti-PODXL antibody were connected via elongated collecting ducts marked with anti-CALB1 antibody, similar to the collecting duct tree observed in the renal system (Figure 3A,B). Both kidney organoids differentiated under normal and hypoxic conditions contained CALB1+/CDH1+ collecting duct cells on day 21; however, the length of a continuous collecting duct was greater in the hypoxia-enhanced organoids than in the control organoids (Figure 3B). Flow cytometric analysis revealed a significant increase in CALB1+ cell population under hypoxia (10.40%) compared to that under normoxia (4.64%), indicating a higher population of collecting duct cells under the hypoxic condition (Figure 3C and Figure S3A). Similarly, the mRNA level of *CALB1* was significantly upregulated in hypoxia-stimulated kidney organoids (Figure 3E).

**Figure. 3.**
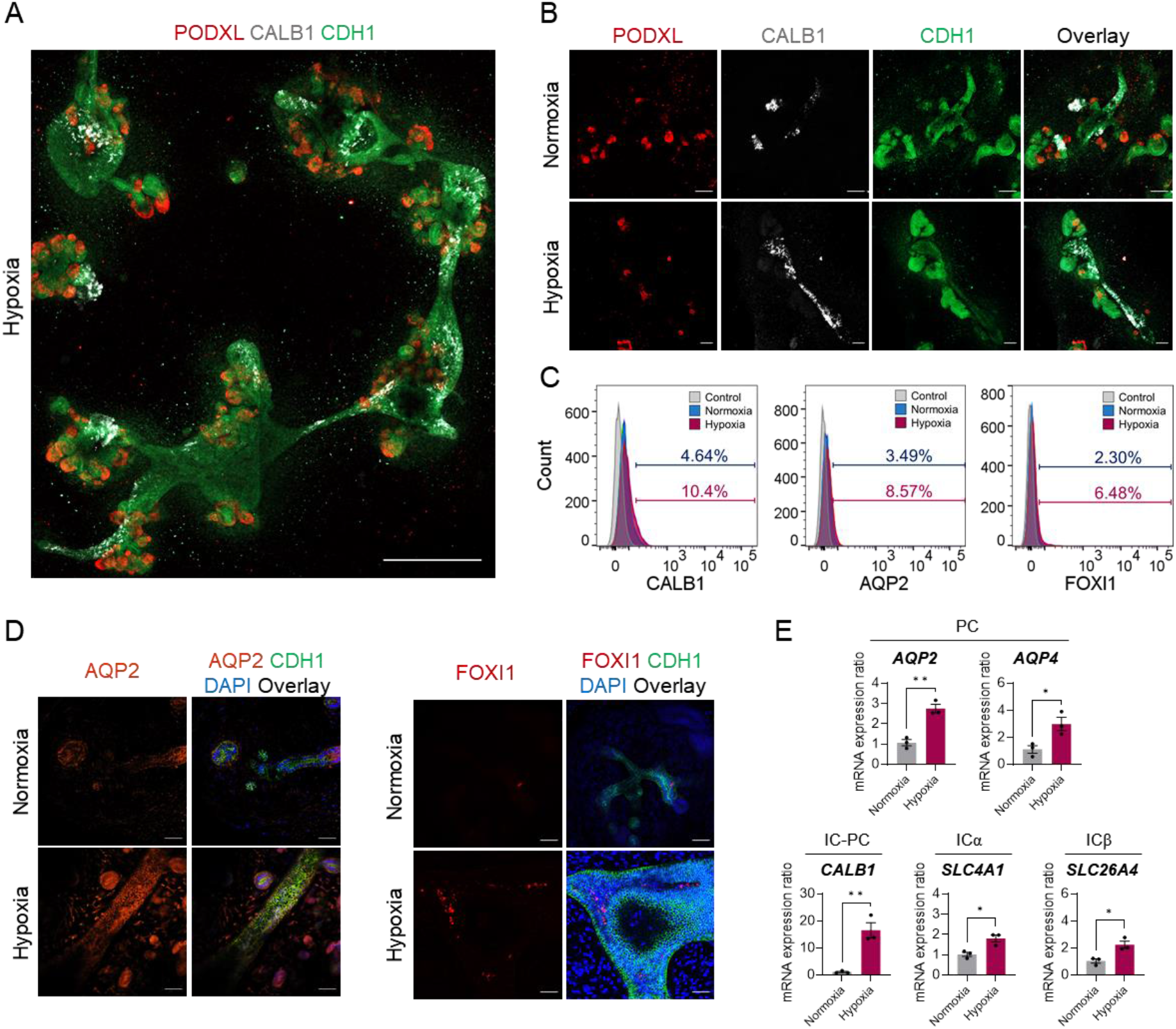
Self-organized kidney organoids spontaneously forming kidney arcades, PCs, and ICs. (**A**) Immunocytochemistry of kidney organoids on day 21 labelled with markers of podocytes (PODXL), distal tubule/collecting duct (CDH1), and collecting duct (CALB1) in collecting ducts showing multiple nephrons connected by collecting duct tree, which is similar to the *in vivo* kidney structure when differentiated under hypoxic conditions. Scale bars, 200 µm. PCs, principal cells; ICs, intercalated cells. (**B**) Immunofluorescent images of kidney organoids on day 21 labelled with PODXL, CDH1, and CALB1, demonstrating higher expression of CALB1 under hypoxic conditions than under normoxic conditions. Scale bars, 100 µm. (**C**) Flow cytometric analysis showing representative frequencies of CALB1+, AQP2+, and FOXI1+ cells in kidney organoids on day 21. Unstained cells were used as controls. (**D**) Immunocytochemistry for AQP2 or FOXI1 as markers for PCs or ICs, respectively, and for CDH1 in kidney organoids on day 26, differentiated under normoxic and hypoxic conditions. Scale bars, 100 µm. (**E**) qRT-PCR analysis of genes encoding markers of each cell type; PCs (*AQP2* and *AQP4*), ICs-PC (*CALB1*), ICα (*SLC4A1*), and ICβ (*SLC26A4*) in kidney organoids on day 26, differentiated under hypoxic and normoxic conditions. PC, principal cells; ICα, alpha intercalated cells; ICβ, beta intercalated cells. Statistical analysis is two-tailed unpaired t-test (\**P* < 0.05; \*\**P* < 0.001)

The functions of renal collecting duct system are carried out by two major cell populations, principal cells (PCs) and intercalated cells (ICs), which are intermingled throughout the entire network of the collecting duct. PCs concentrate urine and regulate Na+/K+ homeostasis, whereas ICs regulate normal acid-base homeostasis via H+ or HCO_3_-in urine.[33, 34] Immunostaining of kidney organoids with anti-AQP2 and anti-FOXI1 antibodies revealed significantly higher expressions of AQP2 and FOXI1 in duct-like structures in the hypoxia-enhanced organoids than in the normoxia group (Figure 3D). When analyzing AQP2+ and FOXI1+ cell populations using flow cytometry, we observed significant increases in both PCs (8.57% versus 3.49%) and ICs (6.48% versus 2.30%) in hypoxia-enhanced kidney organoids (Figure 3C and Figure S3B,C). Additionally, mRNA expression of *AQP2*, *AQP4*, *SLC4A1*, and *SLC26A4*, which are markers of PCs and ICs, was significantly higher in organoids differentiated under the hypoxic condition than under the normoxic condition (Figure 3E). These findings indicate that hypoxic stimulation successfully improved tubular structures and spontaneously induced PCs and ICs populations.

### Hypoxia-enhanced kidney organoids exhibit *in vivo*-like microstructure

The development of cilia in nephrons and collecting ducts has been linked to kidney development, with average length of cilia in developing kidney being < 1 μm[35] and 3 to 10 μm in quiescent and fully differentiated cells.[36] Electron microscopic analysis revealed that a significantly lower percentage of cells in hypoxia-enhanced kidney organoids (1.6%) had primary cilia that were shorter than 1 μm compared to those of normal kidney organoids (8.3%) (Figure 4A,B). This indicates that hypoxia-enhanced kidney organoids contained relatively few immature cells. In addition, 95% cilia with lengths between 3–10 μm were observed in hypoxia-enhanced kidney organoids, which was 1.5-fold higher than those in normal kidney organoids (Figure 4B), further supporting the fact that the hypoxia protocol increased the maturity of kidney organoids.

**Figure 4.**
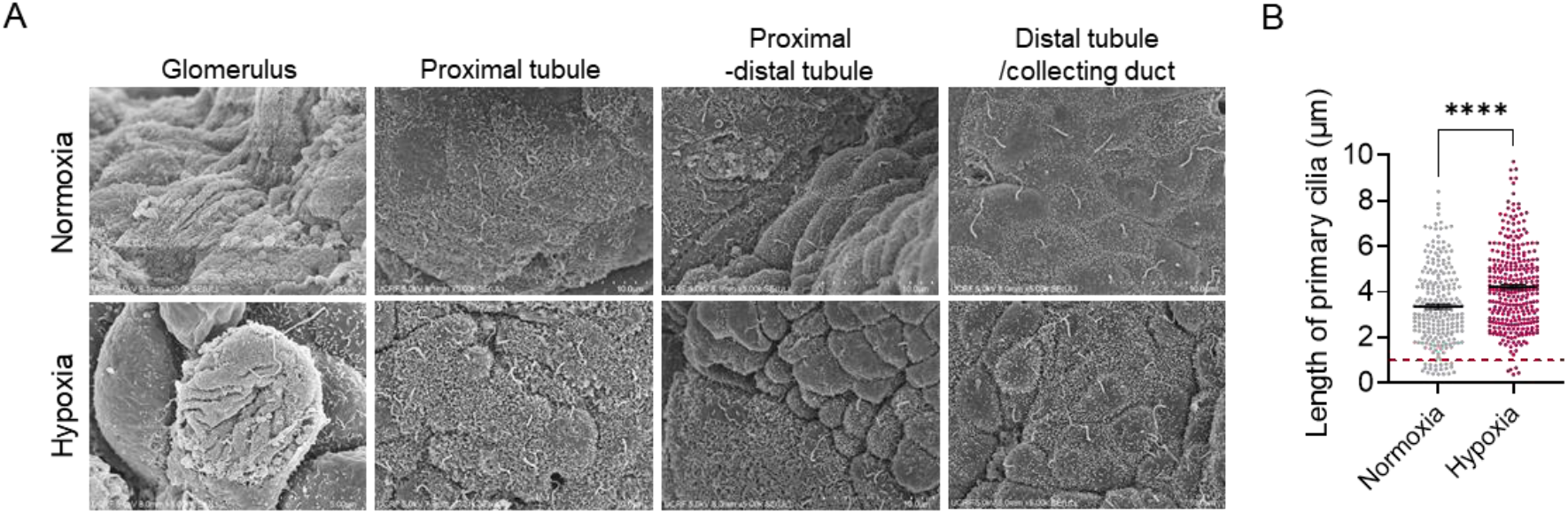
Compartmentalized segments of nephrons in kidney organoids exhibiting matured microstructural features. (A) Electron micrograph of each nephron segment of human kidney organoids differentiated under normoxic and hypoxic conditions. (B) Lengths of primary cilia in kidney organoids on day 21, analyzed by ImageJ. All data are plotted as mean ± S.E. and n = 3 for the independent experiments. Statistical analysis is two-tailed unpaired t-test (\*\*\*\**P* < 0.00001).

Furthermore, the proximal tubules had a well-developed brush border that increases its apical surface area (Figure 4A). We observed denser and longer microvilli in the proximal tubules of hypoxia-enhanced kidney organoids than in normal kidney organoids (Figure 4A). Distinctive short and sparse microvilli were also observed in the distal tubules and collecting ducts of hypoxia-enhanced organoids, which were similar to that of mammalian renal tissue,[37] whereas the distal tubules and collecting ducts in normoxic kidney organoids exhibited relatively flatter structures owing to less developed microvilli (Figure 4A). Both organoids exhibited glomerular podocyte-like structures with numerous projections (Figure 4A). These findings confirm that developmentally-inspired hypoxic stimuli supported the maturation of tubular epithelia in kidney organoids.

### Enhanced reciprocal signaling interactions between MM and UB

To understand the cause of hypoxic stimuli allowing the generation of kidney organoids with higher-order structures, we investigated the mutual induction between UB and MM, which is essential for forming complex kidney structures *in vivo*.[2, 3] UB stimulates MM to differentiate into glomeruli and renal tubular epithelia, and in turn, MM furthers the branching of UB and helps it differentiate into collecting ducts. In mice, UB outgrowth from IM is observed around embryonic day 10.5 owing to the effect of glial cell line-derived neurotrophic factor (GDNF) acting on UB epithelia through its receptor Ret.[38] FGF10 works with GDNF to encourage UB growth.[39] Unexpectedly, we did not find significant upregulation of *GDNF* mRNA levels in the hypoxia group (Figure 5A). As GDNF is activated by SIX2,[40] the decreased expression of *SIX2* on day 9 may lead to reduced *GDNF* expression on day 14 (Figure 1H and Figure 5A). In contrast, we detected *FGF10* mRNA expression only in the hypoxia group (Figure 5B), suggesting that increased *FGF10* expression may have played a role in enhancing UB outgrowth.[39] In response, UB secretes WNT9b that activates WNT4 and promotes the differentiation of some NPCs into nephron components.[41–43] We discovered that *WNT9B* mRNA levels increased by the hypoxia-stimulation protocol (Figure 5A), possibly owing to UB stimulation; likewise, MM expression significantly upregulated *WNT4*, which is similar to the *in vivo* developmental process (Figure 5A).[43] WNT11 in UB tips creates a positive feedback loop in the GDNF/Ret pathway, thereby contributing to the formation of collecting duct system.[44, 45] Compared to the normoxia group, *WNT11* mRNA expression was significantly elevated on day 21 in the hypoxia group (Figure 5A), indicating an improved signal of collecting duct maturation in hypoxia-stimulated kidney organoids.

**Figure 5.**
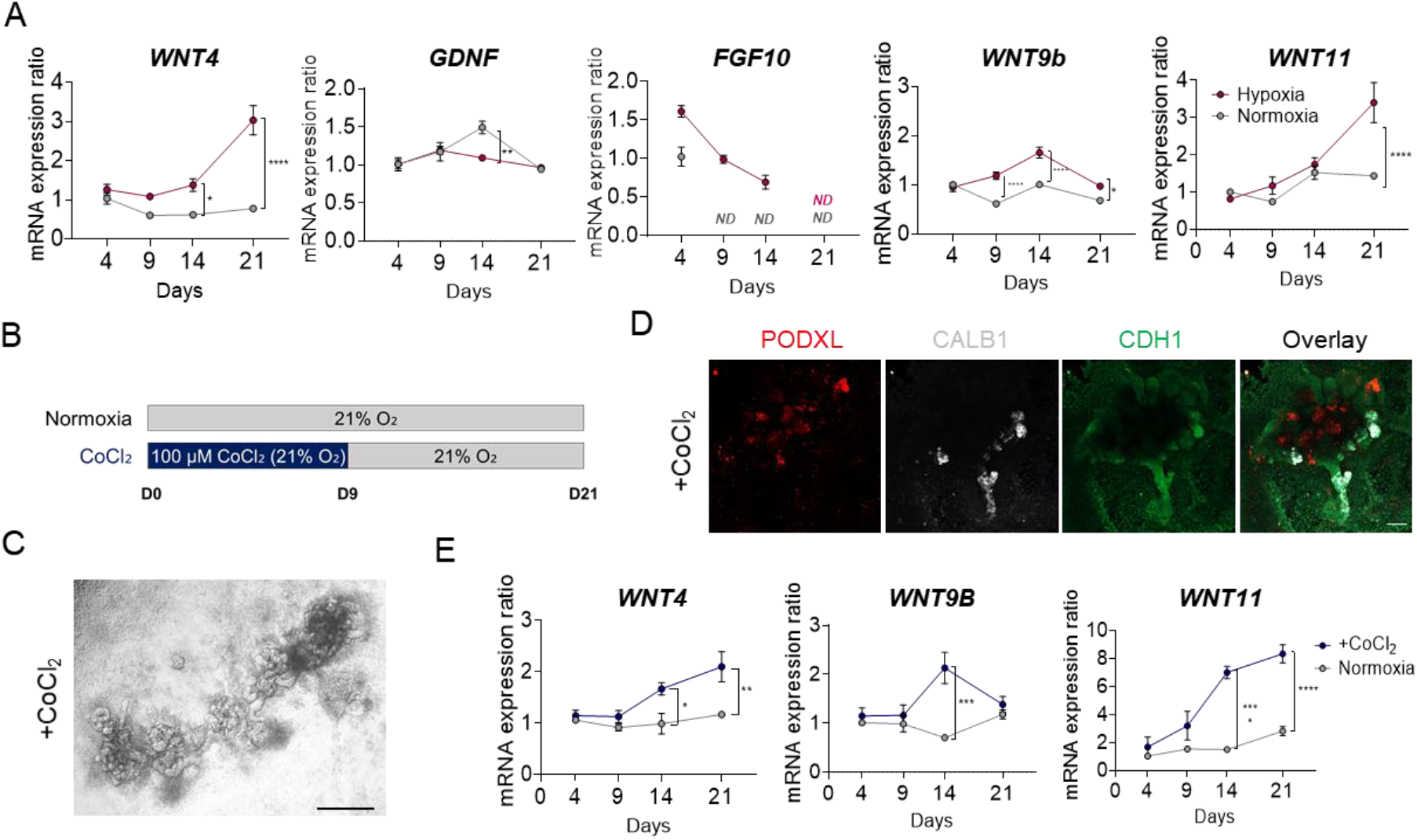
Improved branching morphogenesis by HIF1α-induced commutual inductive signals. (A) mRNA expression ratios of genes encoding *WNT4*, *GDNF*, *FGF10*, *WNT9B*, and *WNT11* analyzed over time and compared to their expression on day 4 under hypoxic versus normoxic conditions. ND; non-detected (B) Timeline for *in vitro* differentiation of kidney organoids using CoCl_2_ as a hypoxia-mimetic agent. (C) Bright-field images of CoCl_2_-treated kidney organoids on day 21. Scale bar, 500 µm. (D) Immunofluorescence micrograph of kidney organoids treated with CoCl_2_ for initial nine days, labelled with antibodies against PODXL, CALB1, and CDH1. Scale bar, 100 μm. (E) mRNA expression ratios of genes encoding *WNT4*, *WNT9B*, and *WNT11*, which are involved in reciprocal interactions between UB and MM compared to their expression on day 4 under hypoxic versus normoxic conditions. All data are plotted as mean ± S.E. and n = 3 for the independent experiments. Statistical analysis is two-way ANOVA followed by Šídák’s multiple comparison test (\**P* < 0.05; \*\**P* < 0.001; \*\*\**P* < 0.0001; \*\*\*\**P* < 0.00001).

Previous studies have indicated that the Wnt/β-catenin signaling pathway interacts with HIF1α signaling during development.[46–48] We noticed increased levels of HIF1α under hypoxic conditions using western blot and confocal immunofluorescent microscopic analyses (Figure 1F,G). To determine whether HIF1α induces the mutual interaction between MM and UB, we treated hiPSCs with 100 µM CoCl_2_ under normal oxygen levels for initial nine days (Figure 5B); CoCl_2_ imitates hypoxia by stabilizing HIF1α[49]. Both CoCl_2_-treated and hypoxia-stimulated kidney organoids had similar structures with multiple nephrons arranged around the UB-derived collecting duct tree (Figure 5C,D). Interestingly, the expression of *WNT4*, *WNT9B*, and *WNT11* was upregulated in CoCl_2_-treated kidney organoids (Figure 5E), exhibiting patterns which are similar to those observed in hypoxia-stimulated kidney organoids. These findings suggest that the accumulation of HIF1α under 5% O_2_ for the initial nine days upregulated WNT signaling, promoting mutual interactions between UB and MM and resulting in enhanced branching morphogenesis, which were similar to the features of normal kidney development.

### Transcriptome analysis of hypoxia-enhanced kidney organoids

To investigate the differentiation process of kidney organoids, we analyzed bulk RNA expression by comparing the kidney organoids on day 21 generated by original protocol under the normoxic condition and our hypoxia-enhanced protocols and identified 1124 differentially expressed genes (DEGs) with a fold change > 2 and p-value < 0.05 (Figure 6A, B). Of these, 560 genes were upregulated and 564 were downregulated in the hypoxia-enhanced kidney organoids compared to those in the normoxic ones (Figure S4A), indicating substantial changes in gene expression owing to different oxygen levels. A generated heatmap for genes encoding markers of each kidney segment revealed decreased genes that are enriched in NPCs and early nephrons, whereas many genes involved in the function of podocytes, proximal tubules, distal tubules and collecting ducts were upregulated in kidney organoids differentiated under the hypoxic condition compared to those under the normoxic condition (Figure 6A).

**Figure. 6.**
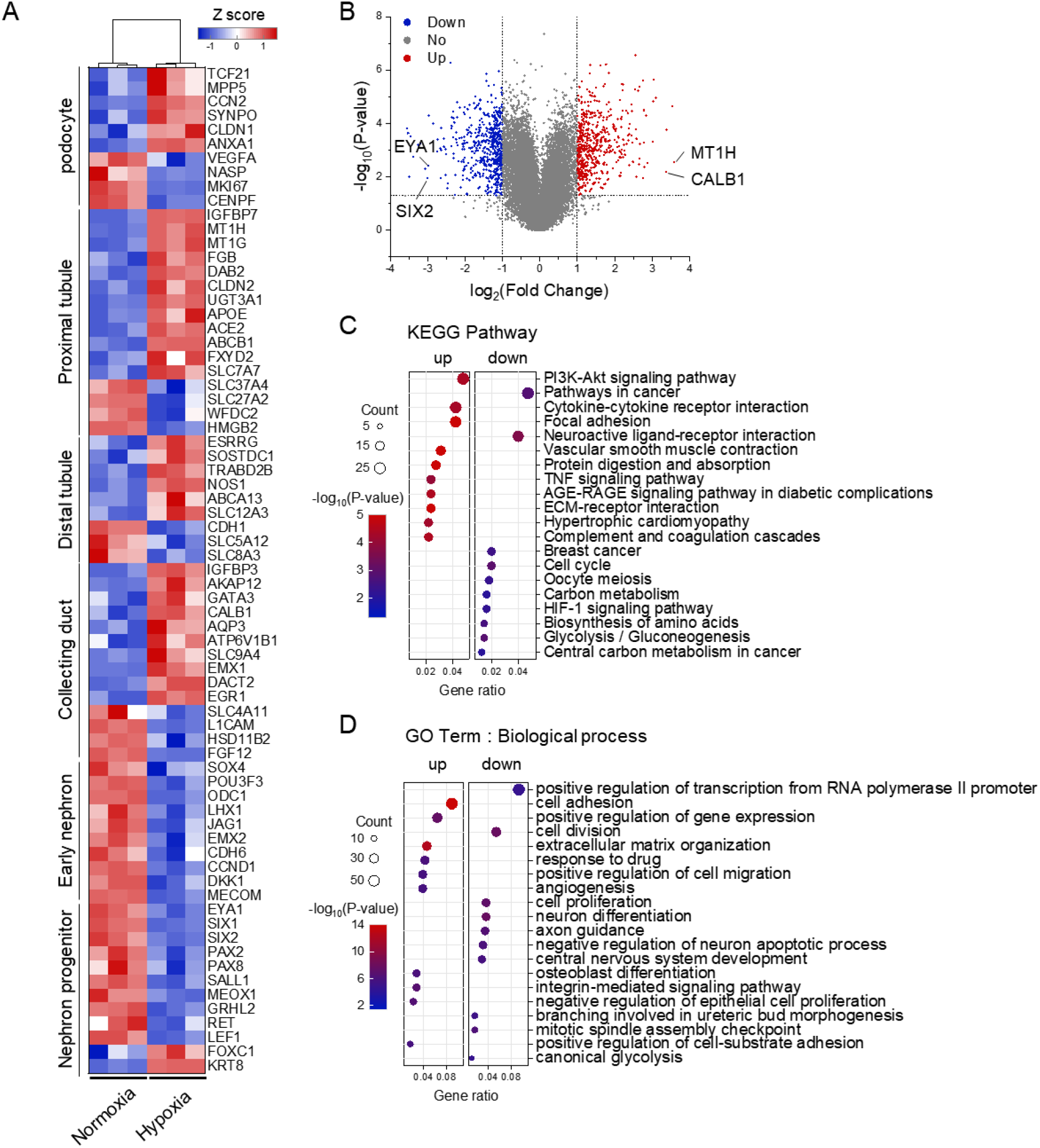
Transcriptomic analysis of kidney organoids differentiated under hypoxic and normoxic conditions. (A) Heatmap and hierarchical clustering of the selected differentially expressed genes (DEGs) expressed in each segment of kidney with fold change > 2.0 and p < 0.05. (B) Volcano plot of the DEGs in kidney organoids on day 21. Genes differentially expressed with fold change > 2.0 and p < 0.05 were marked in red or blue. (C and D) KEGG functional classification (C) and biological process GO term (D) of kidney organoids. The colour of dots represents the rich factor, while the size represents the input number of genes for each KEGG (C) and GO (D) terms. The horizontal axis indicates the significance of enrichment. All data are from 3 independent experiments.

Among the 10 top upregulated genes were *MT1H* and *CALB1* that were enriched in mature proximal tubular and collecting duct cells, respectively (Figure 6B), indicating that tubular maturation was enhanced by the hypoxia protocol. Conversely, *SIX2* and *EYA1* that are required for expansion of NPC, were among the 10 top downregulated genes (Figure 6B). Consistent with this, mature markers of the nephron segments and collecting duct were positively regulated, whereas early nephron and renal progenitor markers were negatively regulated under hypoxia compared to those under normoxia (Figure 6A, B). According to the results of KEGG pathway enrichment analysis, the PI3K–Akt signaling pathway that is involved in normal UB branching during kidney development,[50] was most enriched (Figure 6C). Additionally, focal adhesion-related genes, including *ACTN1*, *COL1A1*, *COL6A6*, and *FN1* were highly upregulated (Figure 6C), which is consistent with their roles in kidney morphogenesis[51] and relatively complex tubular structures observed in hypoxia-enhanced kidney organoids. Notably, the HIF-1 signaling pathway-related genes were downregulated in kidney organoids on day 21 after reoxygenation from day 9, followed by culture under the hypoxic condition (Figure 6C).

The most enriched biological process gene ontology (GO) term pertained to cell adhesion and organization of extracellular matrix (Figure 6D). Cell adhesion plays a critical role in the kidney tubular function and morphogenesis.[52] Among the upregulated genes involved in cell adhesion are *CLDN2*[53] and *CLDN11*, which encode tight junction proteins that selectively control the paracellular transport of molecules in the kidney tubules.[54] These findings were consistent with the well-developed tubular structures observed in kidney organoids subjected to hypoxia. Meanwhile, genes associated with the positive regulation of transcription from RNA polymerase II promoter and cell division were downregulated, which was consistent with the suppression of oncogenesis observed with the hypoxia protocol in the KEGG pathway analysis (Figure 6C, D). The enriched molecular function GO term showed an increase in the expression of genes, such as *RYR2*, *CALB1*, *ANXA1*, and *SPARC*, involved in calcium binding, in hypoxia-enhanced kidney organoids (Figure S4B). This observation suggests that the hypoxia protocol may induce active calcium signaling, which is crucial for converting UB and MM into functional nephron epithelium during renal development.[55]

### Hypoxia-enhanced kidney organoids as renal cystic models for effective drug screening

PKDs are a group of inherited disorders that cause the formation of bilateral cysts in the kidneys.[56, 57] Studies on human cyst epithelial cells and PKD animal models have highlighted the crucial role of cAMP in the development of PKD. A decrease in intracellular Ca^2+^ level triggers a change in the cellular response to cAMP, and the activation of extracellular signal-regulated kinase by cAMP leads to cyst growth.[58] Additionally, cAMP stimulates the secretion of chloride and fluid through chloride channels located in the proximal tubules and collecting ducts, including the cystic fibrosis transmembrane conductance regulator (CFTR) and calcium-dependent chloride channel, anoctamin1 (ANO1).[59, 60] Therefore, PKD treatment currently focuses on reducing renal cAMP levels, increasing Ca2^+^, and inhibiting CFTR/ANO1 to decrease cyst growth.[61]

We found that the mRNA levels of *CFTR* and *ANO1* were significantly increased in kidney organoids subjected to hypoxic stimulation, inducing the formation of well-structured tubules (Figure 7B). Based on this finding, we hypothesized that renal cystic tissues could be more effectively modelled by treating hypoxia-stimulated organoids with forskolin that is used to increase cAMP levels in PKD models.[8, 11–15, 62] To test this hypothesis, we monitored cystogenesis in kidney organoids after exposure to forskolin (10 and 30 μM) on day 21 (Figure 7A). Remarkably, multiple renal cysts formed 24 and 48 h after exposure to forskolin in hypoxia-enhanced kidney organoids, whereas normoxic organoids showed less distinct cystogenesis under the same forskolin treatment conditions (Figure 7C). Confocal immunofluorescence confirmed cyst formation in the inner parts of organoids both in normoxia and hypoxia (Figure S5A). Strikingly, cysts were robustly induced in CDH1+ distal tubules under the hypoxic condition (Figure S5A). We further observed a significant dose-dependent increase in the organoids area, resulting from cyst formation by the hypoxia protocol (Figure 7C, D). Additionally, the number of cysts per unit area was significantly higher in hypoxic than in normoxic kidney organoids after exposure to 30 μM forskolin for 48 h (Figure S5B). In patients with PKD, cAMP activates AQP2, resulting in enhanced water permeability of the collecting ducts.[63, 64] Similarly, we found that mRNA expression of *AQP2* significantly increased when hypoxia-enhanced kidney organoids were exposed to 30 μM forskolin (Figure S5C). These results suggest that the hypoxia induction method can efficiently mimic cAMP-mediated cyst formation, even without genetic mutations in *PKD1* or *PKD2*, likely because of robust formation of renal tubules with well-developed ion channels and transporters.

**Figure. 7.**
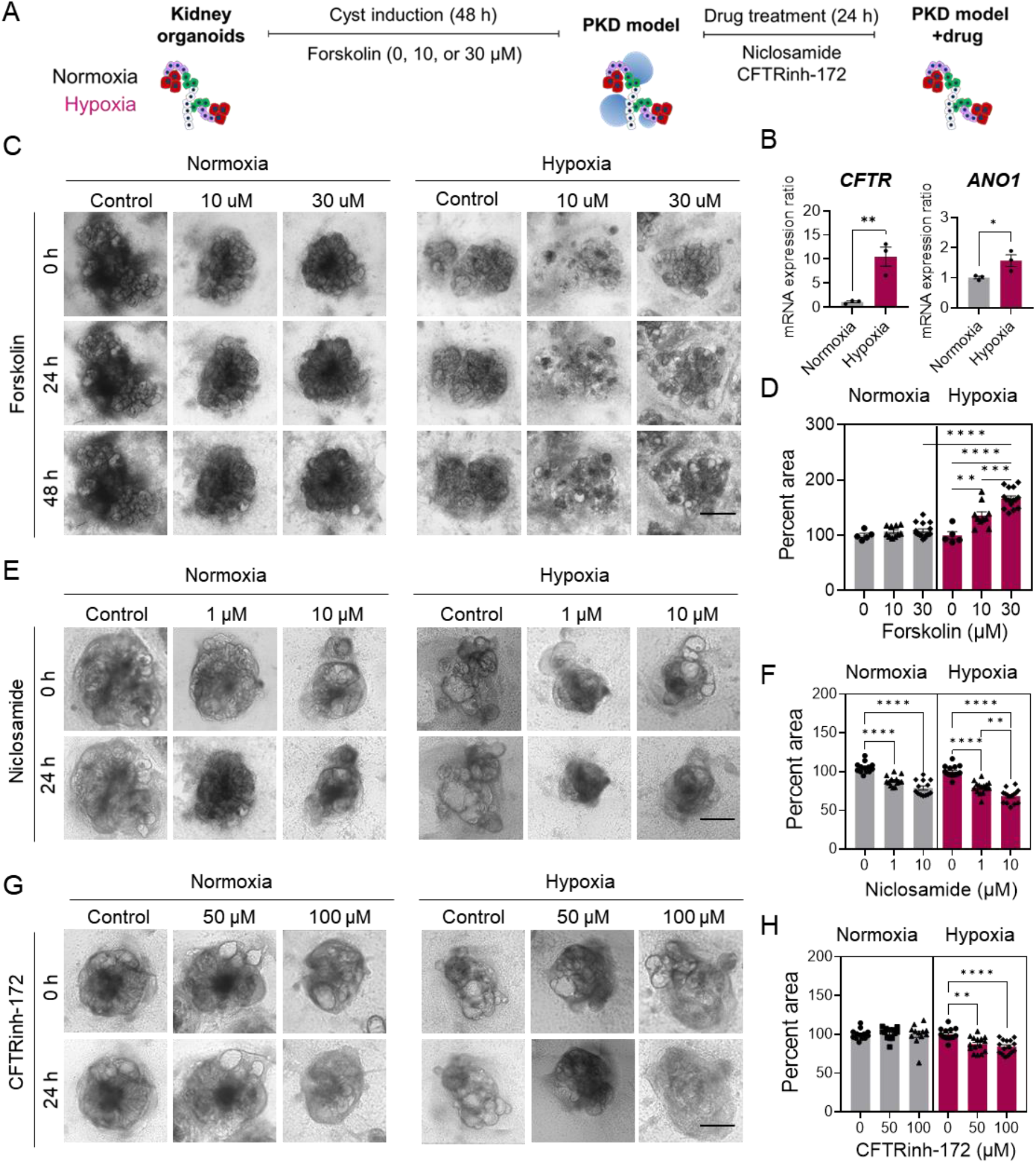
Modelling renal cyst formation with hypoxia-enhanced kidney organoids for reliable screening of PKD drugs. (A) Schematic of the timeline for cyst induction in kidney organoids and drug treatment. (B) Relative mRNA expression of genes encoding chloride channels (*CFTR* and *ANO1*) in kidney organoids differentiated under hypoxic and normoxic conditions. *P* values were determined by two-tailed unpaired t-test (\**P* < 0.05; \*\**P* < 0.001). (C) Bright-field images of forskolin-treated kidney organoids differentiated under normoxic and hypoxic conditions. Scale bars, 500 µm. (D) Percent of area of organoids treated with forskolin compared to that before induction of cystogenesis. (E and G) Bright-field images of cystic kidney organoids treated with chloride channel inhibitors, niclosamine and CFTR inhibitor 172 (CFTRinh-172) for 24 h. The cystic kidney organoids were generated by treating organoids with forskolin 30 µM for 48 h. Scale bars, 500 µm. (F and H) Percent of area of kidney organoids after treatment with (F) niclosamine or (G) CFTRinh-172 for 24 h compared to that of the non-treated group. Statistical analysis is one-way ANOVA followed by Tukey’s multiple comparison test (D, F and H) (\**P* < 0.05; \*\**P* < 0.001; \*\*\**P* < 0.0001; \*\*\*\**P* < 0.00001). All data are plotted as mean ± S.E. and n = 3 for the independent experiments.

To evaluate the potential of our kidney model as a drug-screening tool for PKDs, we tested the therapeutic effects of inhibiting ANO1 and CFTR in the renal cystic model. The ANO1 inhibitor, niclosamide approved by the Food and Drug Administration, effectively suppressed cyst growth by blocking Cl^−^ current and inhibiting ANO1 expression.[65] Cyst-induced kidney organoids treated with forskolin for 48 h were treated with niclosamide for an additional 24 h, and cyst growth was monitored by microscopy (Figure 7A). Both normoxia and hypoxia-stimulated kidney organoids showed a decrease in cyst size when treated with 1 and 10 μM niclosamide, resulting in a significant reduction in organoids size (Figure 7E, F) CFTR inhibitor 172 (CFTRinh-172), an allosteric inhibitor that targets the cytoplasmic face of CFTR, slows cyst growth in PKD mice and *in vitro* models.[65, 66] Interestingly, a significant decrease in the size of kidney organoids was observed with CFTRinh-127 at 50 and 100 μM, particularly in hypoxia-stimulated organoids with relatively high *CFTR* mRNA expression but not in normoxic kidney organoids (Figure 7E, G). These results demonstrate that the proposed organoids can efficiently reproduce the therapeutic effects of PKD drugs by reducing cysts.

## DISCUSSION

This study demonstrates that a hypoxic microenvironment can be mimicked during kidney organoid generation by lowering oxygen levels or using HIF1α-inducing mimetics (e.g. CoCl_2_), to enable the formation of complex structures containing both NPCs and UB-lineage cells. This is the first study to demonstrate that oxygen regulation enhances the formation of tubular structure in kidney organoids, which is more developed than that observed in previous studies.[4–8] The Morizane protocol that particularly generates NPC-derived cells, was modified by changing oxygen tension. When adapted to a hypoxic environment for the first 9 days, ducts derived from UB connected NPC-derived tubules, recreating the structure of the kidney tissue. The maturity of kidney organoids further increased, as they expressed longer primary cilia and highly expressed the proximal tubules, distal tubules and collecting ducts, probably owing to the reciprocal interactions between MM and UB during differentiation under hypoxia versus normoxia conditions. The medium used in organoid culture may successfully differentiate NPCs without UB; however, this study demonstrates that the maturity of NPC-derived cells can be greatly enhanced when they are generated with UB.

Previous observations of kidney formation during embryological development in a hypoxic environment inspired us to study enhanced differentiation using hypoxia.[26] Previous studies have shown that HIF1α were accumulated in progenitor cells,[26] and when we applied similar conditions to differentiate kidney organoids *in vitro*, they exhibited a similar developmental program. Moreover, the Wnt pathway and *FGF10*, indicating the interaction between MM and UB required for the formation of a complete kidney structure, were also enhanced. To ensure that these changes were associated with HIF1α-mediated pathways, we treated the organoids with a HIF1α-eliciting mimetic, CoCl_2_. On day 21, markers indicative of the proximal tubules, distal tubules, and collecting ducts and Wnt signaling were upregulated. Interestingly, the timing of exposure to hypoxia and the level of oxygen tension had a significant impact on the success of generating kidney organoids. Continuous exposure to 5% oxygen for 14 days or 9 days in 1% oxygen tension did not generate proper kidney structures; however, 9 days of adequate hypoxic stimuli led to improved tubular formation. This implies that the temporal control of hypoxia and well-regulated oxygen tension are essential for successful generation of kidney organoids. We hypothesize that the biological responses to reoxygenation following hypoxic stimuli for 9 days could be an important factor, which needs further investigation.

Previous studies have focused on creating complex kidney structures by combining individually generated organoids of NPC- and UB-lineages, which is a complex process and do not contain vascular structure.[17–19] In contrast, our hypoxia protocol is relatively simple, and enables the generation of two populations in the same culture dish and the maintenance of vascular structures. It also enables us to study how the co-induction of UB and MM in the developmental process affects the formation of complex kidney structures and the related diseases. Our hypoxia approach can be adapted to other protocols for generating kidney organoids[4, 5, 10] by optimizing oxygen tension according to the strategy of each protocol.

Our study confirms that hypoxia-stimulated kidney organoids have potential as *in vitro* models of renal cystic diseases. Renal cysts in PKDs typically arise from the collecting ducts, but also in the other overall tubules.[67]Although previous kidney organoids protocols have generated PKD models that exhibit the pathophysiology of PKD, cysts were frequently induced in proximal tubules. Along with this partially non-physiological results, the lack of a collecting duct system has limited the screening of effective PKD drugs.[13] Therefore, modelling cyst formation in the UB/collecting duct in humans would be beneficial for related studies and planning future treatment strategies. The suggested hypoxia-enhanced kidney model generates renal cysts in the tubules and collecting ducts, which is a crucial feature for a preclinical drug development tool that can be used to discover functional PKD drugs that decrease cysts throughout the entire kidney structure.

Furthermore, our kidney organoids exhibited longer primary cilia along the tubules and collecting ducts, demonstrating enhanced maturity of our model. This feature has advantages in modelling PKD, as ANO1, located in the primary cilia, is abundantly expressed in the distal tubules and collecting ducts and plays an important role in cystogenesis. ANO1, a calcium-activated chloride channel, induces the secretion of cystic fluids, leading to the growth of kidney cysts in a high-cAMP environment. Hypoxia-stimulated kidney organoids showed relatively high expression of *ANO1* and *CFTR*, enabling efficient establishment of a cystic model with chemical induction using forskolin. Primary cilia have a sensory function also, modulating the influx of calcium under shear stress. *PKD1* or *PKD2*, the main causative genes of PKDs, are located in primary cilia. Combining a microfluidic platform with hypoxia-enhanced kidney organoids having mutations in *PKD1* or *PKD2*, which have well-developed primary cilia and collecting duct systems, would greatly improve our understanding of how abnormal mechano-transduction owing to genetic malfunction accelerates cyst formation in PKD.[68]

Although the hypoxic protocol has many advantages, further optimization and characterization are required. For instance, we did not observe the ureter connected to multiple collecting ducts, which is supposed to be differentiated from the UB, implying that filtrate removal, which is important for functional assessment of the kidney, may not be achieved. The use of advanced culture platforms, including three dimensional bioprinting and micro-physiological systems, would be helpful in recapitulating the kidney structure containing an open nephric duct. Generating kidney organoids with enhanced vascularization and discovering the optimal oxygen conditions for vasculogenesis and tubulogenesis[32] would further improve the complexity of kidney organoid models. mRNA sequencing results showed suppressed oncogenesis in hypoxia-enhanced kidney organoids, indicating that kidney organoids may be safe for use in cell replacement therapy. However, further maturation of cells by chemical and mechanical stimuli and in-depth evaluation of the generated collecting duct cells are essential for their application in tissue engineering.

In summary, our study demonstrated that kidney organoids exposed to developmentally-inspired hypoxic differentiation conditions exhibited enhanced functionality compared to those of previous kidney models. Enhanced reciprocal interaction signaling between MM and UB led to the maturation of kidney organoids, longer primary cilia, higher expression of marker proteins in each tubular segment and collecting duct, and higher efficiency in modelling renal cysts in the entire kidney structure. Therefore, these organoids may be useful for modeling physiology and pathophysiology throughout the entire course of branching morphogenesis of the kidney and for drug screening by efficiently recapitulating the *in vivo* environment of the kidney.

## MATERIALS AND METHODS

### hiPSCs maintenance

IMR90-4 hiPSCs (passages 45–60) purchased from WiCell were maintained in mTeSR1 (85850, Stem Cell Technologies) in 6-well cell culture plates (30006, SPL Life Sciences) coated with 1% (v/v) LDEV-free hESC-qualified Geltrex (A1413302, Thermo Fisher Scientific) in an incubator at 37 °C with 5% CO_2_. hiPSCs were passaged using Versene (15040066, Thermo Fisher Scientific) at 1:4 to 1:6 ratio every three–four days according to the manufacturer’s protocol (http://www.wicell.org).

### Differentiation of hiPSCs

Kidney organoids were generated using a previously published protocol[6] with modification of the O_2_ conditions. hiPSCs were dissociated using Accutase (A6964, Sigma-Aldrich) and then seeded at a density of 3.8 × 10⁴ cells per well in 1% Geltrex-coated 24-well cell culture plates (30024, SPL Life Sciences) in mTeSR1 supplemented with Y-27632 dihydrochloride (10 μM) (1254, Tocris Bioscience). The medium was replaced with mTeSR1 the following day. When the cells reached 40– 60% confluence, they were washed with phosphate-buffered saline (PBS) and treated with 8 μM CHIR (SML1046, Sigma-Aldrich) and 10 ng mL^−1^ Noggin (120-10C, Peprotech) in basic differentiation medium composed of advanced RPMI 1640 (12633-020, Thermo Fisher Scientific) and 100× GlutaMAX (35050-061, Thermo Fisher Scientific) for four days. On day 4, 10 ng mL^−1^ Activin A (338-AC, R&D Systems) was added for three days, followed by the addition of 10 ng mL^−1^ FGF9 (100-23, Peprotech) and incubation for seven days. 3 μM CHIR was added from day 9 to 11. From day 14, only basic differentiation medium was given and replaced every three days. When renal vesicle-like morphology was observed on day 10, CHIR was removed, and basic differentiation medium supplemented with 10 ng mL^−1^ FGF9 was used. For hypoxic, cells were transferred into a hypoxic incubator at 37 °C (New Brunswick Galaxy 48 R, Eppendorf) with 1% or 5% O_2_–5% CO_2_–N_2_ balance continuously. To mimic hypoxia using CoCl_2_, cells were exposed to 100 µM CoCl_2_ (232696, Sigma-Aldrich) from days 0 to 9 in an incubator at 37 °C with 5% CO_2_.

### Induction and inhibition of cyst formation

To induce cysts in kidney organoids, the previously described basic differentiation medium was supplemented with 10 or 30 μM forskolin (F-9929, LC Laboratories) every day from day 21. To inhibit cyst formation, 30 μM forskolin was administered on day 21 for 48 h, followed by treatment with either 1 or 10 μM niclosamide ethanolamine (5.33087, Sigma-Aldrich) or 50 or 100 μM CFTRinh-172 (HY-16671, MedChemExpress) together with 30 μM forskolin for 24 h. Organoids were imaged using THUNDER Imaging Systems (Leica Microsystems), and their sizes were measured using ImageJ (NIH).

### Immunofluorescence microscopy

Cells were fixed with 4% paraformaldehyde (PFA) in PBS for 2 h at room temperature or overnight at 4 °C. Fixed cells were washed with PBS and blocked in a blocking buffer [3 vol% fetal bovine serum (FBS), 1 wt% bovine serum albumin (BSA), 0.5 vol% Triton X-100 (T8787, Sigma-Aldrich), and 0.5 vol% Tween 20 (P1379, Sigma-Aldrich) in PBS] for 2 h. The antibodies used for immunostaining are listed in the Table S1. Cells were incubated with primary antibodies diluted in the same buffer for 3 to 5 days at 4 °C, and then washed in PBST [0.1 vol% Triton X-100 in PBS]. Secondary antibodies conjugated with Alexa Fluor-488, 555, 647, or Streptavidin conjugated Cy5 in same buffer were used for staining overnight at 4 °C. Cells were washed with PBST and then incubated for 2 h at room temperature in PBS with 2 μg mL^−1^ DAPI (D9542, Sigma-Aldrich). Cells were imaged using an LSM980 confocal microscope (Zeiss). ImageJ was used to compare the colocalization of markers and area of the nephron segments.

### qRT-PCR

mRNA expression was analyzed using qRT-PCR. An AccuPrep Universal RNA Extraction Kit (K-3140, Bioneer) was used to extract total RNA. For cDNA synthesis, an AccuPower RocketScript Cycle RT PreMix (K-2202, Bioneer) and a T100 Thermal Cycler (Bio-Rad) were used. qRT-PCR was performed using SYBR Green Realtime PCR Master Mix (QPK-201, TOYOBO) on a CFX Connect Real-Time PCR Detection System (Bio-Rad). All samples were run in two technical replicates and normalized with respect to GAPDH levels using the 2^-ΔΔCt^ method. The primer sequences are listed in Supplementary Table S2.

### Bulk RNA sequencing

Total RNA was isolated from kidney organoids on day 21 using TRIzol. Sequencing libraries were prepared using a TruSeq Stranded mRNA Library Prep Kit (Illumina). Sequencing was performed using a NovaSeq 6000 system (Illumina). Sequence reads were extracted in FASTQ format and trimmed from both ends based on a minimum read length of 36 bp. Reads were mapped to the human genome hg38 using HISAT2 v.2.1.0., and transcripts per kilobase million (TPM) were obtained using StringTie v.1.2.3b. Volcano plots and heat maps were generated using normalized gene expression represented by log_2_(TPM+1) and z-scores, respectively. KEGG pathway and GO term enrichment analyses were performed using the DAVID database. Plots were generated using OriginPro.

### Western blot

Cells were lysed in 1 M Tris-HCl, pH 6.8, 8 wt% sodium dodecyl sulphate (SDS), 40 vol% glycerol containing beta-mercaptoethanol, and bromophenol blue and separated on an SDS–polyacrylamide gel. Proteins in the gel were transferred onto a polyvinylidene fluoride membrane (88520, Thermo Fisher Scientific) and then blocked with 3 wt% BSA. The blots were incubated in PBS containing 0.1% Tween-20 and 3% BSA with anti-HIF1α antibody (1:1000) (AF1935, Novus Biologicals) or anti-beta actin antibody (1:1000) (ab8227, Abcam) and then incubated with HRP-conjugated anti-rabbit (1:3000) (65-6120, Thermo Fisher Scientific) and anti-goat (1:5000) (sc-2020, Santa Cruz Biotechnology) antibodies. Protein bands were visualized using a Pierce ECL Western Blotting Substrate (32106, Thermo Fisher Scientific).

### Scanning electron microscopy

Organoids were fixed using 2.5% glutaraldehyde in 0.1 M sodium cacodylate buffer for 1 h and washed for 2 min in PBS. Samples were further fixed with 1% osmium tetroxide (75633, Sigma-Aldrich) for 1 h and washed with distilled water. Serial dehydration was performed in 25%, 50%, 75%, and 95% ethanol solutions and absolute ethanol for 3 min. After dehydration, hexamethyldisilazane (440191, Sigma-Aldrich) was used. The samples were coated with platinum using sputter (Hitachi) and imaged using SU8220 (Hitachi). ImageJ was used to measure the lengths of primary cilia.

### Dextran uptake assay

Kidney organoids were incubated in basic differentiation medium with 100 μg mL^−1^ of 10,000-MW cascade blue-labelled dextran (D1976, Thermo Fisher Scientific) for 24 h and then cultured in fresh basic differentiation medium for 24 h. Organoids were fixed with 4% PFA and stained with LTL.

### Flow cytometry

Kidney organoids were detached by gentle pipetting. Harvested organoids were singularized using Accutase (A6964, Sigma-Aldrich) for 20 min, and dissociated cells were fixed with 4% PFA for 30 min at room temperature. Cells were permeabilized with permeabilization buffer [0.2 vol% Triton X-100 and 5 mM ethylenediaminetetraacetic acid (EDTA) in PBS] for 15 min at room temperature and then blocked with 5% donkey serum (5 vol% donkey serum and 5 mM EDTA in PBS) for 15 min at room temperature. After incubation with anti-CALB1, FOXI1, or AQP2 antibodies (1 μg mL^−1^) in flow cytometry staining buffer (2 vol% FBS, 2 wt% BSA, 5 mM EDTA and 1% sodium azide) for 30 min at 4 °C and washed with the washing buffer. FACSVerse (BD Biosciences) and FlowJo (BD Biosciences) were used for flow cytometry analysis.

### Statistical analysis

All data represent mean ± standard error (S.E.). Statistical analysis was performed using Student’s t-test, one-way analysis of variance (ANOVA) with Tukey’s multiple comparison test, or two-way ANOVA with Šídák’s multiple comparison test using GraphPad Prism v.9; \**P* < 0.05; \*\**P* < 0.001; \*\*\**P* < 0.0001; \*\*\*\**P* < 0.00001.

## Acknowledgements

This study was supported by a National Research Foundation of Korea (NRF) grant funded by the Ministry of Science and ICT (NRF-2022M3A91015716, 2021R1A4A3030597, 2019M3A9H1103769, and RS-2023-0020870) and a Research Fund (1.230039.01) of Ulsan National Institute of Science and Technology.

## Author contributions

H.L. designed the study, conducted overall data acquisition, analysis, and visualization with T.-E.P. and D.S.K., who supervised all work. D.K. contributed to analyze the data. H.Y. contributed in western blot analysis. J.H.K contributed in scanning electron microscopy. H.L. and T.-E.P. prepared the manuscript with input from all others.

## Data and materials availability

All data needed to evaluate the conclusions in this paper are present in the paper and/or the Supporting information.

