## Supplementary Information for "A developmentally-inspired hypoxia condition promotes kidney organoid differentiation from human pluripotent stem cells"

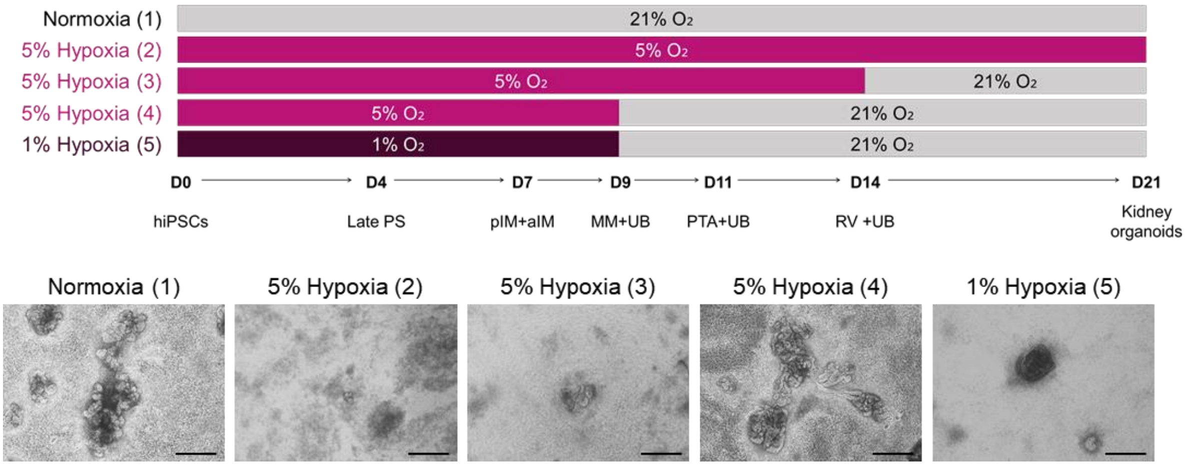


**Figure S1. Differentiated kidney organoids depending on oxygen conditions.** Timelines and corresponding bright field images of kidney organoids differentiated in each oxygen tension condition on day 21. hPSCs, human induced pluripotent stem cells; PS, primitive streak; pIM, posterior intermediate mesoderm; aIM, anterior intermediate mesoderm; MM, metanephric mesenchyme; UB, ureteric bud; PTA, pre-tubular aggregate; RV, renal vesicle.

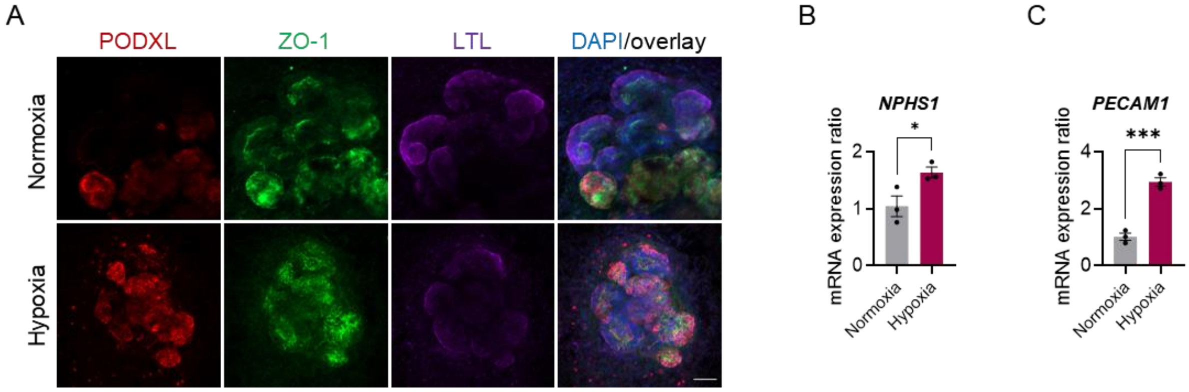


**Figure S2. Matured features of podocytes in hypoxia**. (**A**) Immunocytochemistry for markers of podocytes (PODXL), tight junction (ZO-1) and proximal tubules (LTL) on day 21 of differentiation under normoxia and hypoxia conditions. Scale bars = 100 µm. **(B-C)** mRNA expression ratios of glomerular filtration barrier (*NPHS1*) and endothelial cell (*PECAM1*) in kidney organoids on day 21 differentiated under hypoxia versus normoxia conditions. All data are plotted as mean ± S.E. and n = 3 for the independent experiments. Statistical test: Two-tailed unpaired t-test (**P* < 0.05; ****P* < 0.0001).

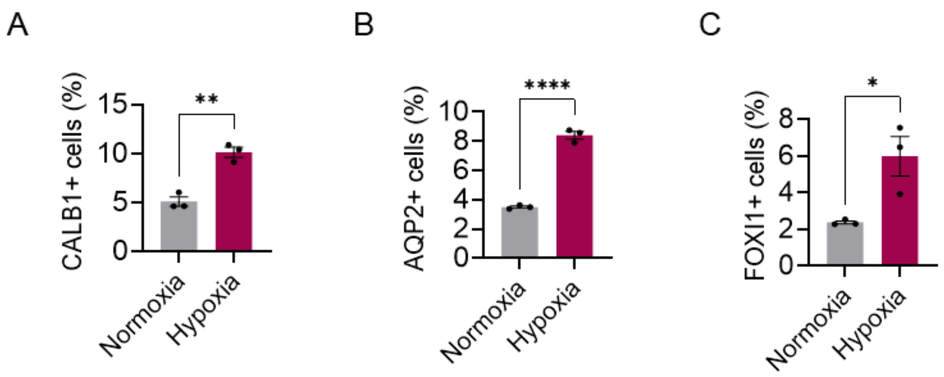


**Figure S3 Hypoxic effects on collecting duct cells.** (**A-C**) Flow cytometry analysis showing that CALB1+, AQP2+, and FOXI1+ cells on day 21, and 30 kidney organoids under hypoxia compared to normoxia. All data are plotted as mean ± S.E. and n = 3 for the independent experiments. Statistical test: Two-tailed unpaired t-test. (**P* < 0.05; ***P* < 0.001; *****P* < 0.00001).

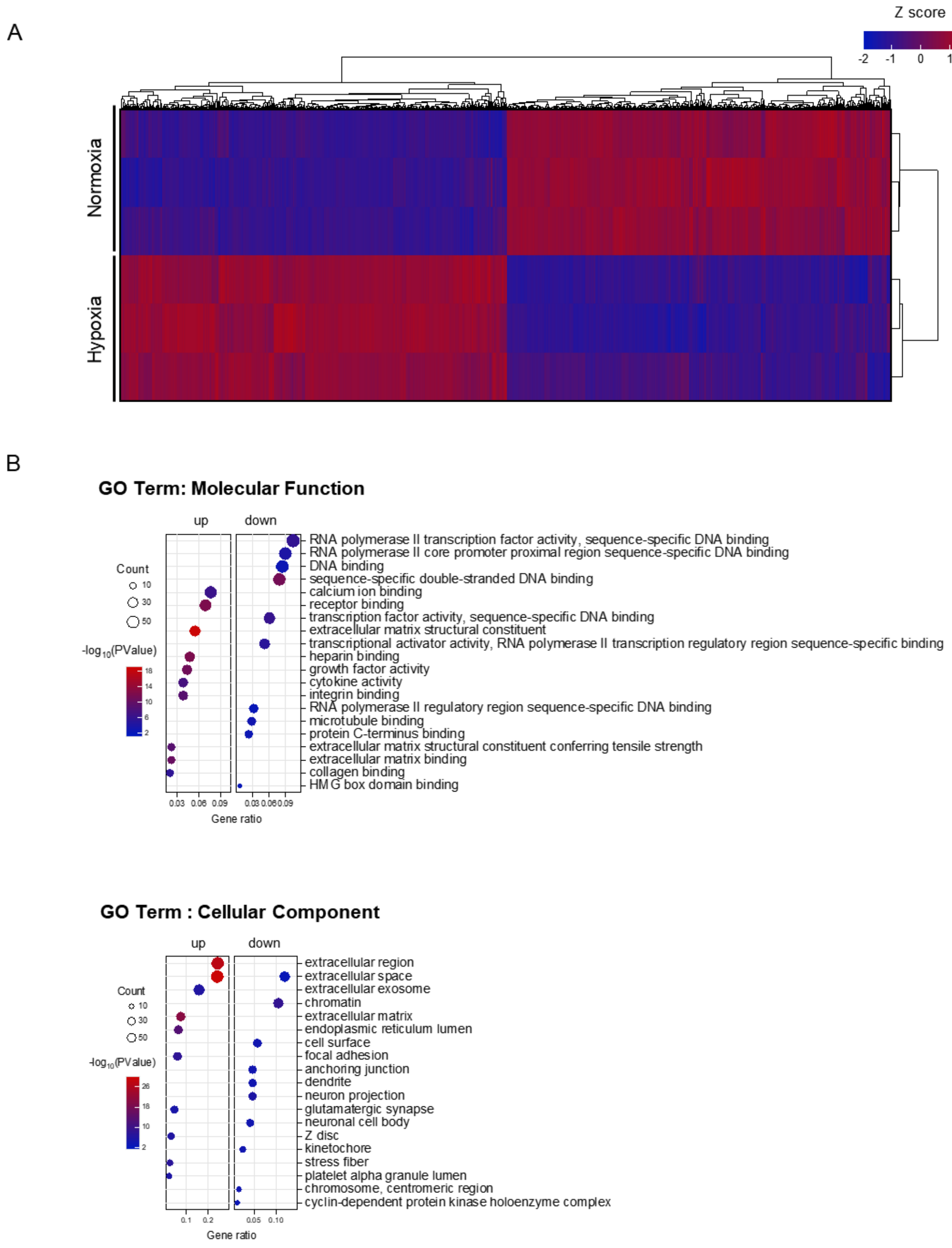


**Figure S4 Differentially expressed genes in kidney organoids under hypoxia versus normoxia conditions.** (**A**) Heatmap of DEGs with fold change over 2.0 and $p < 0.05$ in hypoxic and normoxic conditions. (**B**) Molecular function and cellular component GO Terms of the kidney organoids under hypoxia compared to normoxia. GO Terms on Y axis were arranged by descending gene ratio on X axis. The size and color gradient of the dots represent significance and affected number of genes. All data are from 3 independent experiments.

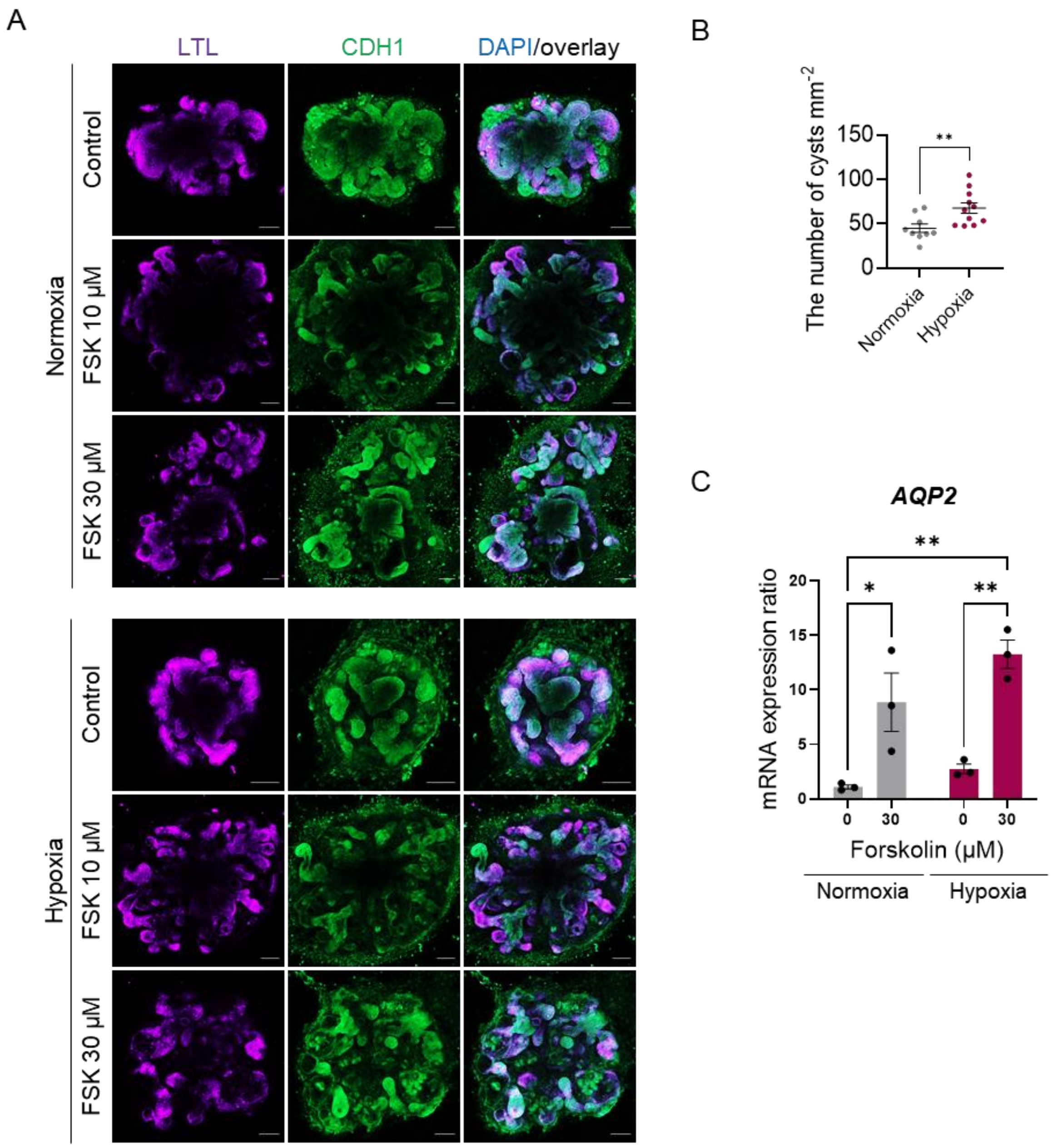


**Figure S5. Validation of cyst induced PKD models in hypoxic and normoxic conditions.** (**A**) Immunocytochemistry for a marker of proximal tubules (LTL) and distal tubules/collecting ducts (ECAD) in hypoxia and normoxia. (**B**) The number of counted cysts per mm2 depending on treatment of 30 μM forskolin in normoxic and hypoxic kidney organoids compared to non-treated conditions in normoxia. Statistical analysis is two-tailed unpaired t-test (***P* < 0.001). (**C**) qRT-PCR showing the expression of *AQP2* in kidney organoids cultured under hypoxic versus normoxic conditions when treated with forskolin 30 μM. Statistical analysis is one-way ANOVA followed by Tukey's multiple comparison test (*$P < 0.05$; **$P < 0.001$). All data are plotted as mean ± S.E. and n = 3 for the independent experiments.

**Table S1.** List of antibodies for immunocytochemistry

| Antibody | Source | Catalog # | Concentration |
|---|---|---|---|
| AQP2 | Novus Biologicals | NB110-74682 | 1:100 |
| AQP4 | Thermo Fisher Scientific | PA5-53234 | 1:200 |
| CALB1 | Sigma-Aldrich | C9848 | 1:100 |
| PECAM1 | abcam | ab9498 | 1:100 |
| CDH1 | Proteintech | 20874-1-AP | 1:100 |
| CDH1 | abcam | ab11512 | 1:300 |
| FOXI1 | Novus Biologicals | NB300-926 | 1:300 |
| HIF1α | Novus Biologicals | AF1935 | 1:50 |
| LHX1 | Developmental Studies Hybridoma Bank | 4F2-c | 1:500 |
| LTL | Vector Laboratories | B-1325 | 1:200 |
| PAX2 | Novus Biologicals | AF3364 | 1:200 |
| PODXL | R&D Systems | AF1658 | 1:200 |
| SIX2 | Proteintech | 11562-1-AP | 1:500 |
| TBX6 | R&D Systems | AF4744 | 1:100 |
| UMOD | abcam | ab167678 | 1:100 |
| WT1 | Novus Biologicals | NBP2-67587 | 1:200 |
| ZO-1 | Thermo Fisher Scientific | 40-2200 | 1:100 |
| Donkey Anti-Goat IgG H&L (Alexa Fluor® 555) | abcam | ab150134 | 1:200 |
| Donkey Anti-Goat IgG H&L (Alexa Fluor® 647) | abcam | ab150135 | 1:200 |
| Donkey Anti-Mouse IgG H&L (Alexa Fluor® 488) | abcam | ab150109 | 1:200 |
| Donkey Anti-Rabbit IgG H&L (Alexa Fluor® 488) | abcam | ab150061 | 1:200 |
| Donkey Anti-Rabbit IgG H&L (Alexa Fluor® 647) | abcam | ab150063 | 1:200 |
| Streptavidin, Cy5 | Vector Laboratories | SA-1500-1 | 1:200 |

**Table S2.** List of primer sequences for qRT-PCR

| Gene | Forward | Reverse |
|---|---|---|
| *ABCB1* | GGGAGCTTAACACCCGACTTA | GCCAAAATCACAAGGGTTAGCTT |
| *ANO1* | CTGATGCCGAGTGCAAGTATG | AGGGCCTCTTGTGATGGTACA |
| *AQP1* | TAACCCTGCTCGGTCCTTTG | AGTCGTAGATGAGTACAGCCAG |
| *AQP2* | CTCCCTCCTCTACAACTACGTG | CTCCTCCCAATCGGTGTCC |
| *AQP4* | CATGGAAATCTTACCGCTGGT | TCAGTCCGTTTGGAATCACAG |
| *TBXT* | CTGGGTACTCCCAATGGGG | GGTTGGAGAATTGTTCCGATGA |
| *CALB1* | TCCAGGGAATCAAAATGTGTGG | GCACAGATCCTTCAGTAAAGCA |
| *CFTR* | TGCCCTTCGGCGATGTTTTT | GTTATCCGGGTCATAGGAAGCTA |
| *CDH1* | ATTTTTCCCTCGACACCCGAT | TCCCAGGCGTAGACCAAGA |
| *FGF10* | CAGTAGAAATCGGAGTTGTTGCC | TGAGCCATAGAGTTTCCCCTTC |
| *GAPDH* | TGTGGGCATCAATGGATTTGG | ACACCATGTATTCCGGGTCAAT |
| *GDNF* | GGCAGTGCTTCCTAGAAGAGA | AAGACACAACCCCGGTTTTTG |
| *NPSH1* | GGCTCCCAGCAGAAACTCTT | CACAGACCAGCAACTGCCTA |
| *PECAM1* | AACAGTGTTGACATGAAGAGCC | TGTAAAACAGCACGTCATCCTT |
| *SIX2* | GGCCAAGGAAAGGGAGAACA | GAGCTGCCTAACACCGACTT |
| *SLC26A4* | TGGTGGGATCTGTTGTTCTGA | GGATCTGCCAAGTACCTCACT |
| *SLC34A1* | TCACGAAGCTCATCATCCAG | TTCCTCAGGGACTCATCACC |
| *SLC4A1* | GGTGATGGACGAAAAGAACCA | AAGACTCTACGCAGCTCTAGG |
| *UMOD* | ATGTGGGGCCAATGACATGAA | CAGTCCCGGTTGTCTCTGT |
| *WNT4* | AGGAGGAGACGTGCGAGAAA | CGAGTCCATGACTTCCAGGT |
| *WNT9b* | TGTGCGGTGACAACCTCAAG | ACAGGAGCCTGATACGCCAT |
| *WNT11* | ATGTGCGGACAACCTCAGCTAC | GATGGAGCAGGAGCCAGACA |
